# Honey bee *Apis mellifera* L. Responses to Oxidative Stress Induced by Pharmacological and Pesticide Compounds

**DOI:** 10.1101/2023.08.11.553037

**Authors:** Faizan Tahir, Michael Goblirsch, John Adamczyk, Shahid Karim, Mohamed Alburaki

## Abstract

The western honey bee, *Apis mellifera* L., is a eusocial insect that plays major roles in ecosystem balances and pollination of plants and food crops. Honey bees face multiple biotic and abiotic stressors, such as pathogens, diseases, chemical pesticides, and climate change, which all contribute to honey bee colony loss. This study investigated the impacts of multiple pharmacological and pesticide molecules on honey bee survival and gene regulation responses. In an 11-day cage experiment, sublethal doses of tunicamycin, thapsigargin, metformin, paraquat, hydrogen peroxide, and imidacloprid were administered to newly emerged sister bees. Daily treatment consumption and mortality were recorded, as well as the transcription expression of twelve major genes (*AChE-2*, *Apisimin*, *Apidaecin*, *mrjp1*, *Sodq*, *cp450*, *SelT*, *SelK*, *Ire1*, *Xbp1*, *Derl-1*, *Hsc70*), some of which are markers of oxidative and endoplasmic reticulum (ER) stresses in honey bees. At day 9 of the treatments, protein damage was quantified in caged bees. Kaplan-Meier model indicated significant (*p* < 0.001) toxicological effects of paraquat, H_2_O_2_ and tunicamycin on bee survivorship compared to controls with better survivals for other molecules. Post-ingestive aversion responses were recorded only in the case of tunicamycin, hydrogen peroxide and imidacloprid. Nonetheless, significantly higher protein damage on day 9 was only identified in bees exposed to paraquat and imidacloprid. Some antioxidant genes significantly regulated vis-à-vis specific treatments. Our results reveal age-related regulation of other major genes with significant inter-gene positive correlations.

## INTRODUCTION

The western honeybee, *Apis mellifera* L., is a eusocial insect that is arguably the most important pollinator insect, which also provides important contributions to human life through its different hive products and pollination of food crops. Honey bees, including other non-*Apis* bees, pollinate 80% of the world’s flowering plants and more than ninety different food crops (Rader *et al*. 2016) with a total estimated crop value of $17 billion annually in the United States (Calderone 2012). Besides their pollination services and agricultural importance, honey bees are ecologically vital in maintaining balanced ecosystems (Klein *et al*. 2007; Potts *et al*. 2010). Honey bee colonies face an assorted number of biotic and abiotic stressors which include, but are not limited to diseases, parasites, pathogens, and chemical pesticides, which all have led to significant losses during the last two decades (Vanengelsdorp *et al*. 2011; Steinhauer *et al*. 2014; Kulhanek *et al*. 2017; Li *et al*. 2018). Honey bee colony loss is a complex problem to tackle as such phenomenon seems to be multifactorial and is the product of synergistic effects between various bee stressors (Alaux *et al*. 2010; Vidau *et al*. 2011; Aufauvre *et al*. 2012; Boncristiani *et al*. 2012; Doublet *et al*. 2015).

Honey bee biological processes and responses to stressors have been investigated in laboratory cage experiments, which provide a more controlled environment and limit the implication of other variables on the outcomes (Evans *et al*. 2009; Gregorc *et al*. 2018; Alburaki *et al*. 2019a; Alburaki *et al*. 2022). Previous studies have shown that nutritional stress and pathogenic infections (i.e., Deformed Wing Virus, *Nosema ceranae*) have a severe impact on honey bee colony strength (Branchiccela *et al*. 2019). The first physiological step in the cascade of events induced by stressors is the manifestation of cellular oxidative stress. This results from a disturbance in the balance between producing reactive oxygen species (ROS), or free radicals, and antioxidant defenses (Di conza and Ho 2020), and any deficiency in antioxidant defenses can cause damage that occurs at various tissue levels of the organism. The second physiological response to stress in the cell is manifested at the endoplasmic reticulum (ER) level. The ER is a large membrane-enclosed cellular organelle found in all eukaryotes (Hirata *et al*. 2021). At this site, membranes are folded, proteins are secreted, lipids and sterols are synthesized, and free calcium is stored (Di conza and Ho 2020). If the stress persists and is not mitigated at the cellular level, an imbalance between the demand for protein folding and the capacity of the ER for protein folding can occur and leads to protein damage (Di conza and Ho 2020). Despite their difference in nature, stressors lead to physiological stress at the cell level, which in turn challenges the organism’s homeostasis.

Honey bee colonies, as superorganisms, can maintain and return to homeostasis despite stressors such as parasite infestations or exposure to pesticides. Such a phenomenon is known as social resilience (Ulgezen *et al*. 2021), which ensures the survival of the honey bee population as a whole, especially in feral colonies. Such attributes, well manifested by feral colonies, could be harnessed and applied to managed colonies which may help limit the constant losses and improve beekeeping management. Aside from managed honey bee populations, anthropogenic stress has been linked to the decline of feral honey bees (Siviter *et al*. 2023). More specifically, in urban landscapes and home backyards, pesticides are applied for pest management control affecting both managed and feral honey bees.

Gene regulation in honey bees operates in a complex manner to alleviate the effect of multiple biotic and abiotic stressors affecting their survival. Some genes have been linked to specific stressors, while others are still under investigation. For instance, major royal jelly protein 1 (*mrjp1*) plays pivotal roles in honey bee nutrition and larvae development (Srisuparbh *et al*. 2003; Li *et al*. 2021), *Xbp1* and *IRE1* are transcription factors that regulate the expression of genes important to the proper functioning of the immune system and in the cellular stress response (Johnston *et al*. 2016; Adames *et al*. 2020). Sodesque (*Sodq*) catalyzes the conversion of superoxide into oxygen and hydrogen peroxide, thus controlling the levels of various reactive oxygen species (Wang *et al*. 2018). Genes well related to both ER stress and redox stress are two selenoproteins T and K. *SelT* has thioredoxin reductase-like oxidoreductase activity, modulates the contraction processes through the regulation of calcium release (Alburaki *et al*. 2019b; Pothion *et al*. 2020), while *SelK* is involved in ER-associated degradation of soluble glycosylated proteins and plays a role in the protection of cells from ER stress-induced apoptosis (Alburaki *et al*. 2019b; Xia *et al*. 2022). *Apismin* and *apideacin* are both hypothesized to play roles in nutrition and pathogen infections and are considered Anti-Microbial Peptides (AMP) genes (Casteels *et al*. 1989; Shen *et al*. 2007). cytochrome P450 (*Cp450*) codes for enzymes that are membrane-bound hemoproteins that play a pivotal role in the detoxification of xenobiotics, cellular metabolism and homeostasis (Zhang *et al*. 2018).

Oxidative stress on caged honey bees was induced in the current study using six different abiotic stressors; two pesticides (imidacloprid and paraquat) and four pharmacological compounds (tunicamycin, thapsigargin, metformin, and hydrogen peroxide). Imidacloprid is a neonicotinoid insecticide highly toxic for bees and widely used in agriculture for pest management. This molecule acts on several types of post-synaptic nicotinic acetylcholine receptors in the insect nervous system, binds to the nicotinic receptor causing failure of the neuron to propagate any signal. Acetylcholinesterase is a cholinergic enzyme primarily found at postsynaptic neuromuscular junctions like muscles and nerves and it immediately hydrolyzes acetylcholine, which is a naturally occurring neurotransmitter, into acetic acid and choline (Dvir *et al*. 2010). The prime role Acetylcholinesterase is to terminate neuronal transmission and signaling between synapses to prevent acetylcholine dispersal and activation of nearby receptors (Smulders *et al*. 2004; Dvir *et al*. 2010). The sustained activation of the receptor results from the inability of acetylcholinesterase to break down the pesticide, an irreversible process that induces excessive ROS production (Dvir *et al*. 2010; Nicodemo *et al*. 2014; Alburaki *et al*. 2019a). Paraquat catalyzes the formation of ROS through accepting electrons from photosystem I and transferring them to molecular oxygen (Kennedy et al., 2021). Tunicamycin inhibits N-linked glycosylation, which induces ER stress (Hirata et al., 2021); thapsigargin is known to inhibit SERCA (sarco endoplasmic reticulum Ca^2+^ ATPase). This sets off Unfolded Protein Response (UPR) and if protein misfolding is not resolved, will induce ER stress (QUYNH DOAN ET AL., 2015). Metformin reduces redox stress but is reported to induce distinct ER stress pathways in cardiomyocytes (PERNICOVA ET AL., 2014). Hydrogen peroxide works by producing destructive hydroxyl free radicals that can attack membrane lipids, DNA, and other essential cell components (Brudzynski 2020).

In this study, we determined the toxicological effects of sublethal doses of pharmacological inducers and pesticides on honey bee as well as the transcriptional response and protein damage caused by oxidative and ER stresses.

## MATERIALS & METHODS

### 1. Laboratory Bioassay Design

This experiment was conducted at the individual honey bee level with six treatments: tunicamycin, thapsigargin, paraquat, hydrogen peroxide, imidacloprid, and metformin; and two controls: control-H_2_O which consisted of 1:1 sugar and water for the first five treatments and control-PBS made of 1:1 sugar syrup with PBS (Phosphate Buffered Saline 1x, pH 7.4) for the metformin treatment as it is not soluble in water and was dissolved in PBS. Eight hundred one-day old sister bees were emerged in an incubator (35°C, RH 55%) and randomly distributed into eight different cages. These cages were specifically designed for feeding experiments and are fully described in (Gregorc *et al*. 2017). Sublethal concentration treatments were chosen based on available toxicological data at the ECOTOX database (US Environmental Protection Agency EPA) as well as previous investigations to enable data comparison. Tunicamycin was administered at 19,600 ppb, thapsigargin: 195 ppb, metformin: 129,000 ppb, paraquat: 1,000 ppb, hydrogen peroxide: 4,000 ppb and imidacloprid at 20 ppb.

The experiment lasted 13 days and consisted of two phases: 1- two-day acclimation period where bees were allowed to adjust to cage conditions (day -2 to day 0 / no data reported) and 2- treatment period, in which bees were investigated for effects of exposure on survival and at the molecular level at two time points during an 11-day exposure period. During the acclimation period, bees in cages were fed 1:1 sugar syrup without addition of treatment. At the start of Phase 2, (i.e., day 0), cages were randomly assigned to the eight treatments, which were administered *ad libitum* using 20 mL syringes through sugar syrup. The sugar syrup consumption was daily recorded by weighing the syringes using a sensitive scale (± 0.01 g) and calculating read differentials. Dead bees were collected daily and counted for each cage. Nine bees from each treatment were sampled at day 5 and 9 for molecular analyses. Bees were frozen on dry ice and immediately stored at -80°C for further molecular analyses.

### 2. RNA Extraction

RNA was extracted from individual bees using whole-body tissue. For each treatment and date of collection, RNA was extracted from three bees using the Trizol extraction method (Chomczynski 1993) with some modifications to the original protocol. Individual whole bee body was put into 1 mL of TRIzol and crushed using sterile pestles. The lysate was pipetted up and down several times to homogenize the bee tissues and then sonicated (Bioruptor Pico,Diagenode) for 10 cycles consisting of a 30 s pulse and 30 s rest at 4°C. Samples were transferred to a rocker for 10 min at room temperature and then centrifuged at ∼15,000 *g* for 10 min at 4°C. Resulting supernatants were transferred to fresh RNase free tubes and incubated for 5 min at room temperature to permit complete dissociation of the nucleoproteins. Two hundred µL of chloroform was added to the tubes and vortexed aggressively. Samples werethen incubated on a rocker at 4°C for 10 min before being centrifuged at 15,000 *g* at 4°C for 20 min. The resulting aqueous phase was transferred to a new tube. Six hundred µL of isopropanol was added to the samples,vortexed, and then incubated for 10 min at -20°C. RNA was precipitated by centrifugation at 15,000 *g* for 15 min at 4°C. The supernatant was discarded, and the RNA pellet was washed with 600 µL of 70% ethanol and was pipetted up and down 3 timesand centrifuged (15,000 *g*) for 5 min at 4°C. The supernatant was discarded, and the pellet was air dried for 5 min and then resuspended in 50 µL molecular grade RNAse free water. RNA extracts were subsequently inspected using a NanoDrop ND 1000 spectrophotometer (Thermo Scientific) to determine the RNA quantity and quality and stored at -80°C.

### 3. Transcriptional analysis

Gene expression of twelve major antioxidant and developmental genes were evaluated from three whole honey bee samples taken from timepoint day 5 and day 9 (Acetylcholinesterase 2, Apisimin, Apidaecin, Major royal jelly protein 1, Sodesque, Cytochrome P450, Selenoprotein T, Selenoprotein K, Inositol-requiring enzyme 1, X-box binding protein, Derlin 1 and Heat shock 70-kDa protein cognate 3). cDNA was produced from total RNA using BioRad iScript Kit following the manufacturer’s protocol. Previously published primers were confirmed on their respective targets and used in this study, Table 1. The twelve target genes were normalized against two housekeeping genes (glyceraldehyde-3-phosphate dehydrogenase and ribosomal protein S5) known for their stability in honey bee tissues (Scharlaken *et al*. 2008; Alburaki *et al*. 2017). All RT-qPCR runs consisted of 3 biological replicates per treatment and time combination and each biological sample was run with three technical replicates per gene using the following cycling protocol: 95°C for 30 s, followed by 40 cycles of 95°C for 5 s and 60°C for 30 s, and a final melting step of 95°C for 10 s and a 0.5°C increment for 5 s from 65°C to 95°C. All qPCR plates were normalized using an inter-plate calibrator ran on each plate and the two housekeeping genes (Table 1). Gene analysis was conducted using relative normalized expression calculated via the ΔΔCt values through the BioRad Maestro Software. Datasets were subsequently transferred to the R environment (Team 2011) for statistical analysis and figure generation.

**Table 1.**
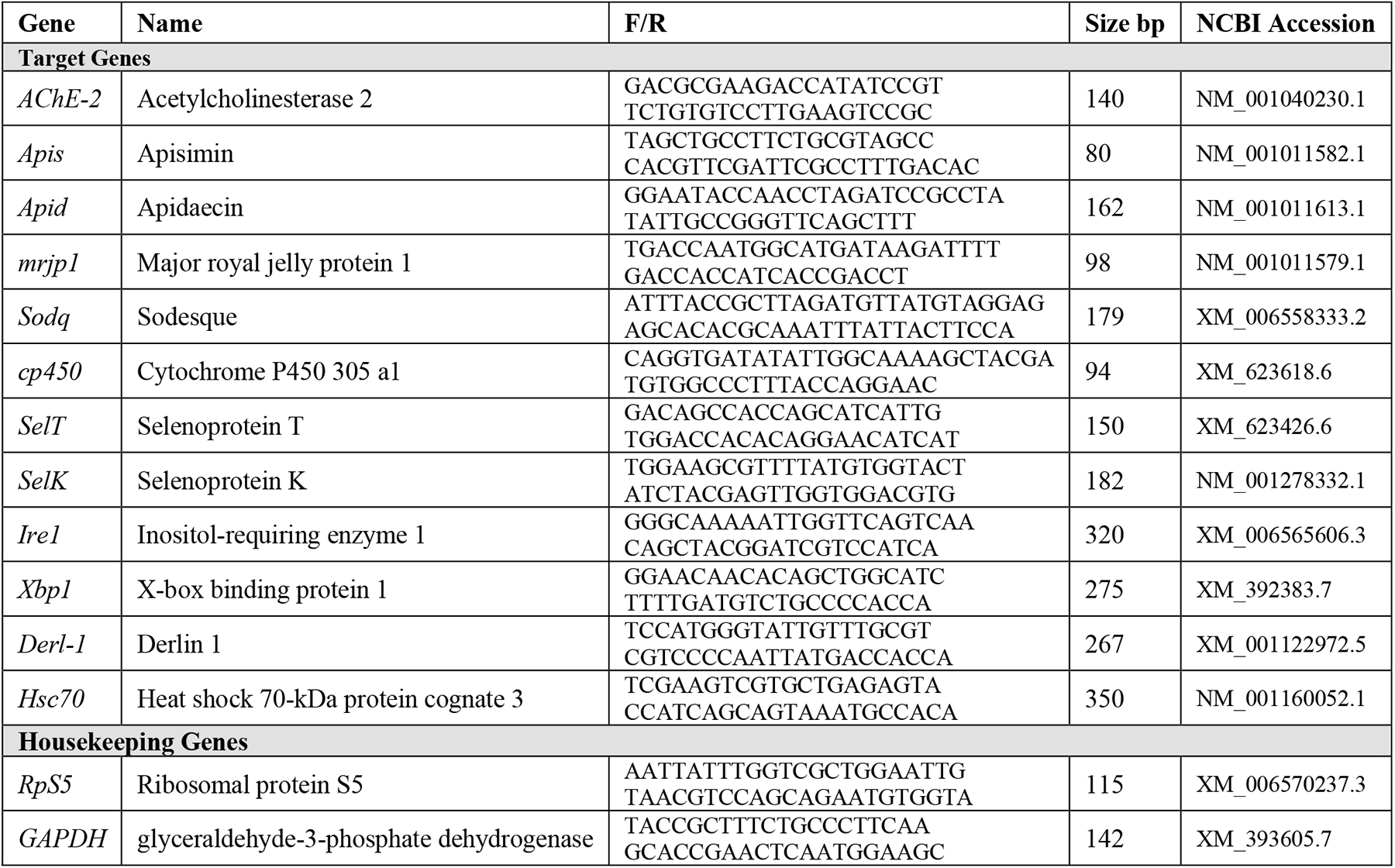
List of the target genes analyzed in this study. Primer sequence for each gene and amplicon size is given as well as the NCBI accession number. Two housekeeping genes known for their stability across honey bee tissues were used to standardize the qPCR dataset.

### 4. Quantification of Protein Damage

To quantify potential post-transcriptional damage caused by the various administrated treatments, a protein carbonyl content assay was conducted on treatments of the last sampling date (Day 9). Protein was extracted from individual bee in triplicate using a protein extraction buffer consisting of 20mM Tris-HCl pH 8.0, 30mM NaCl, and 10% glycerol. The tissues were crushed by using a pestle and sonicated using a Bioruptor Pico (Diagenode) sonication device for 10 cycles of 30s pulse and 30s rest at 4°C. Homogenates were centrifuged at 5,000 *g* for 10 min at 4°C and the supernatants were collected. The protein carbonyl contents in studied samples were estimated using Sigma-Aldrich Kit (MO, USA) as described in the manufacturer’s protocol.

### 5. Statistical Analysis

The cage experiment was conducted at the individual bee level with eight treatments, three biological replicates per treatment and three technical replicates for each gene. All statistical analyses of this study were carried out in the R environment (Team 2011) using RStudio version 2022.12.0+353. First, each dataset was tested for normality using the Shapiro test. Syrup consumption, recorded daily, was calculated at the cage level for each of the eight studied groups. An ANOVA was conducted at a 95% confidential interval on data normality distributed with three levels of significance (P < 0.05*, < 0.001**, < 0.001**). Kruskal–Wallis rank test, a nonparametric test, was used on data that failed the normality test in which multiple comparisons and *p*-values were adjusted with the Benjamini-Hochberg method using the “FSA” Library. H

Survival probability and cumulative hazard were calculated for each treatment group by the Kaplan-Meier survival probability model in R using three Packages: “dplyr”, “survival”, “survminer”. Figures were generated in the same environment utilizing four main libraries: “ggplot2”, “doby”, and “plyr”. Gene regulations were displayed as relative normalized expression per date and overall averages. Heatmaps were generated using the “pheatmap” library either by date (Day 5 and 9) or by overall expression of each studied gene. Correlation in the gene regulations between treatments were performed using the R libraries “PerformanceAnalytics” and “corrplot” with an intermediary level of significance (*p* < 0.01). Principal coordinate analysis PCA was carried out using the “factoextra” library to estimate the expression of each variable on a 3- dimensional scale and treatment group similarity. Box and whisker plots were constructed to visualize the data anddisplay the median, first and third quartiles, and both maximum and minimum values for each condition.

## RESULTS

### 1. Toxicity of Stressors

The Kaplan-Meier model showed significant differences (*p* < 0.001) in survival rates among administered stressors, Fig. 1. The lowest level of mortality during the 11-day experiment was recorded in cages exposed to the Control-PBS treatment while the lowest survival rate was observed in bees exposed to paraquat. Paraquat induced an early mortality starting at day 3 post-administration followed by exposure to hydrogen peroxide (H_2_O_2_), which led to 100% mortality at day 9. Metformin and imidacloprid had a significantly greater survival rate than the control, while tunicamycin led to lower survival rate compared to the control. Caged bees exposed to both thapsigargin, and control had similar survival rates. The cumulative hazard of paraquat was chronic, started early at day 3 but did not lead to total mortality. However, hydrogen peroxide’s cumulative hazard sharply increased at day 8 causing complete mortality of caged bees.

**Figure 1.**
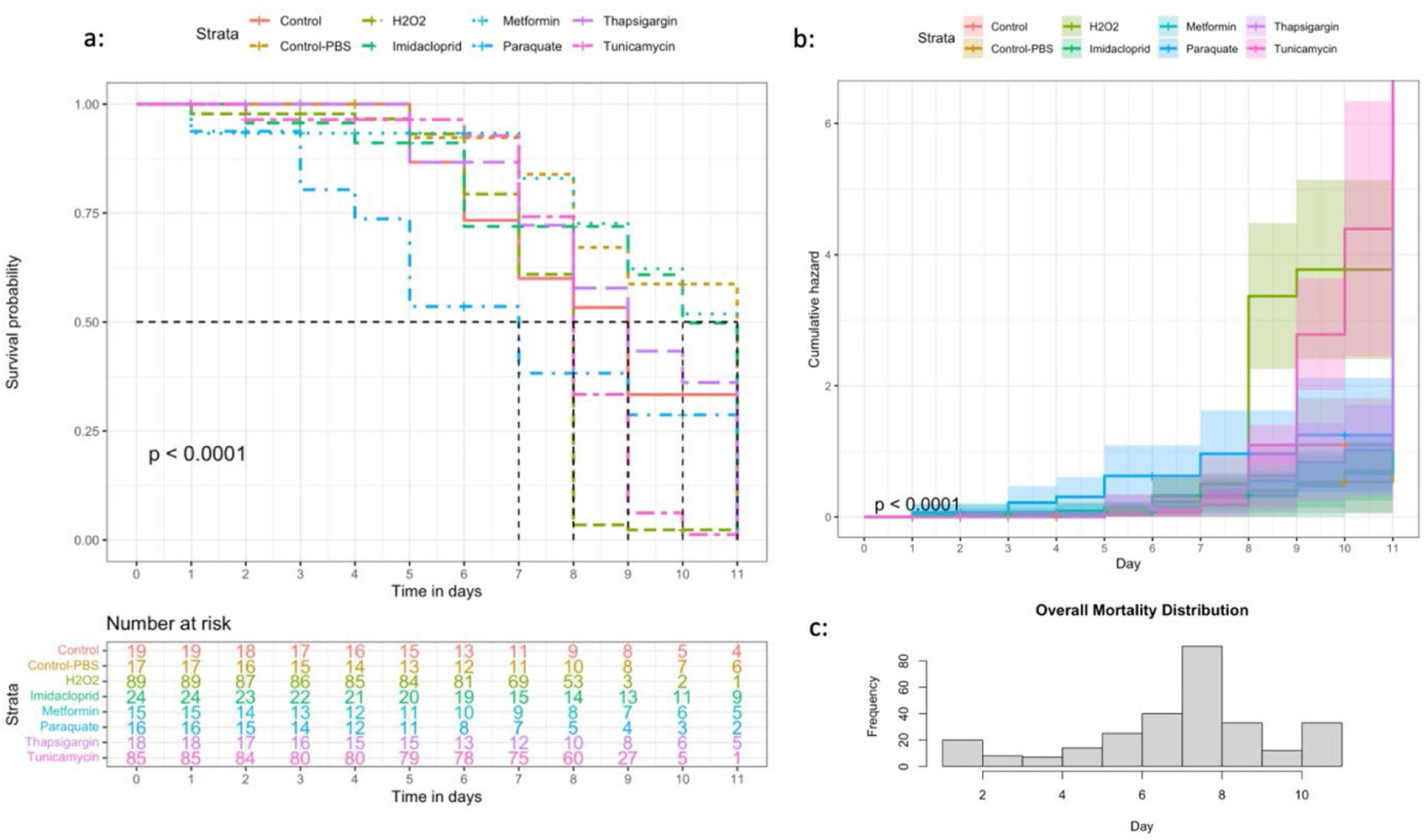
Kaplan-Meier survival probability (a) and cumulative hazard (b) models conducted on the honey bee groups subjected to six different treatments and two controls. The number of honey bees at risk is estimated through the same model for each treatment. Distribution of overall mortality throughout the experiment in day (c). Significant mortality differences among treatments were identified at *p* < 0.001***.

For syrup consumption, caged bees consumed significantly (*p* < 0.001) lower amounts of tunicamycin, H_2_O_2_, and imidacloprid compared to all other treatment compounds including both controls, Fig. 2a. No significant differences were found in the amount of syrup consumed among thapsigargin, metformin, paraquat, and both controlsW. The accumulative consumption graph confirms this finding over the 11-day experiment, Fig. 2b.

**Figure 2.**
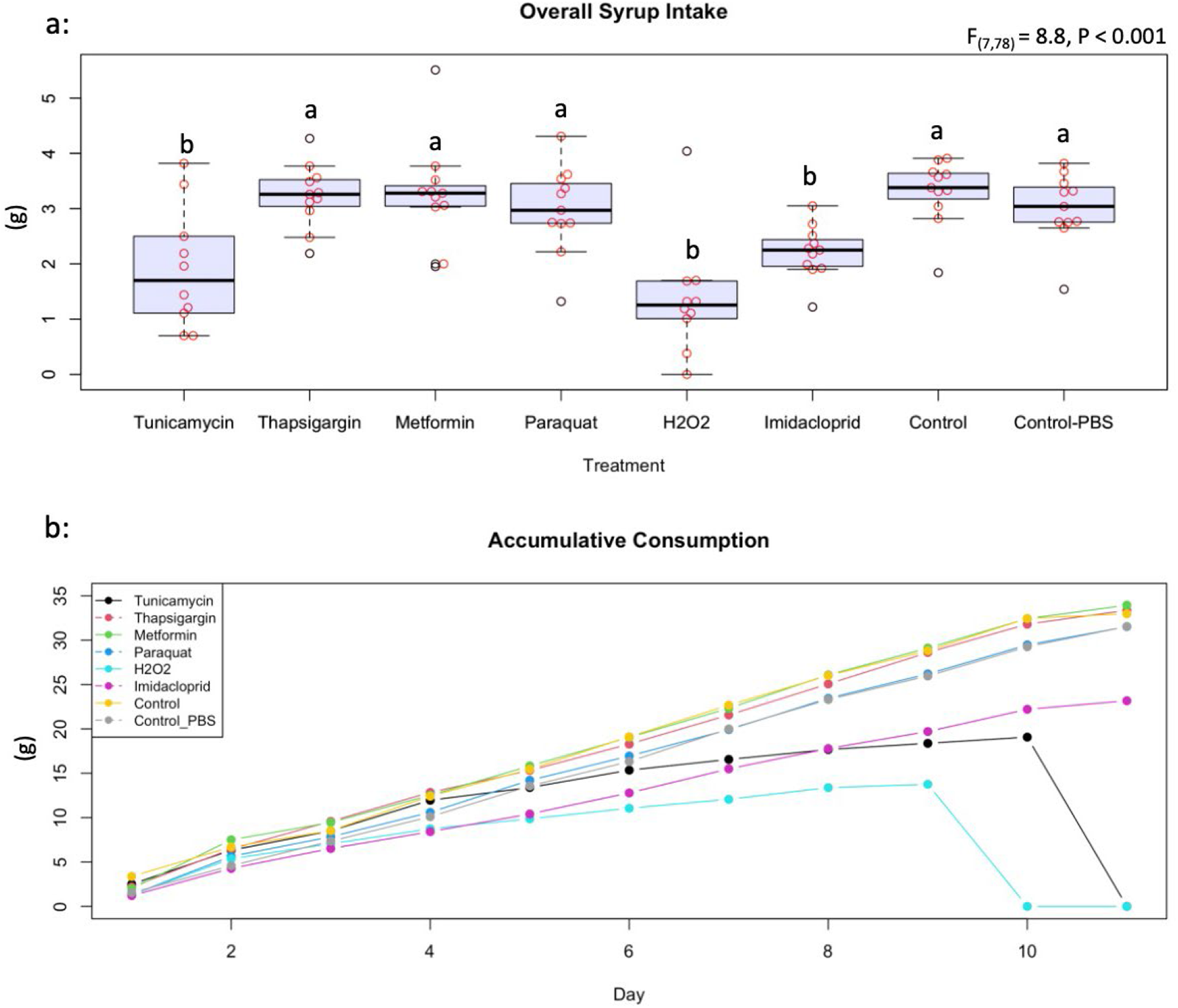
Average *at libitum* syrup intake (a) and accumulative syrup intake (b) displayed by treatment. Accumulative consumptions of the eight administered treatments and controls (tunicamycin, thapsigargin, melformin, paraquat, hydrogen peroxide, imidacloprid, control and control in PBS buffer) are shown in a longitudinal manner. ANOVA was conducted at three levels of significance (*p* < 0.05*, < 0.001**, < 0.001***). Boxplots with different alphabetical letters are statistically significant.

### 2. Transcriptional Analysis and ER Stress

Honey bee response to stresses induced by pharmacological molecules and agricultural pesticides was evaluated by identifying the transcript level of genes involved in the mitigation of oxidative and ER stress. *AChE-2* was significantly (*p* < 0.001) upregulated in day 9 compared to day 5 irrespective of the treatments, while both *apisimin* and *apidaecin* showed no effect of time or treatment on their regulation, except higher (*p* < 0.05) regulation of *apisimin* at day 5 for metformin compared to imidacloprid, Fig. 3. Treatments did not affect the regulation of *mrjp1*, however, an overall significant (*p* < 0.05) drop in its regulation was observed at day 9, Fig. 4. A similar finding was recorded for the regulation of *P450* with significant (*p* < 0.05) upregulation at day 9. Regulation of *Sodq* gene however, was not affected by time, and was significantly higher (*p* < 0.05) for imidacloprid at day 5 compared to tunicamycin, and significantly upregulated for the control compared to tunicamycin at day 9, Fig. 4. Both studied selenoprotein genes (*SelT* and *Selk*) were not affected by the treatments similar to *Ire1,* Fig. 5. Nonetheless, each of these three genes changed regulations vis-à-vis time. *SelT* and *Ire1* were downregulated in day 9, while *SelK* was upregulated in day 9, Fig. 5. *Xbp1* was significantly (*p* < 0.05) upregulated for metformin and thapsigargin compared to imidacloprid at day 5 with a significant (*p* < 0.001) overall downregulation at day 9 irrespective of the treatments, Fig. 6. Regulations of both *Derl-1* and *Hsc70* were not affected by the treatments, Fig. 6.

**Figure 3.**
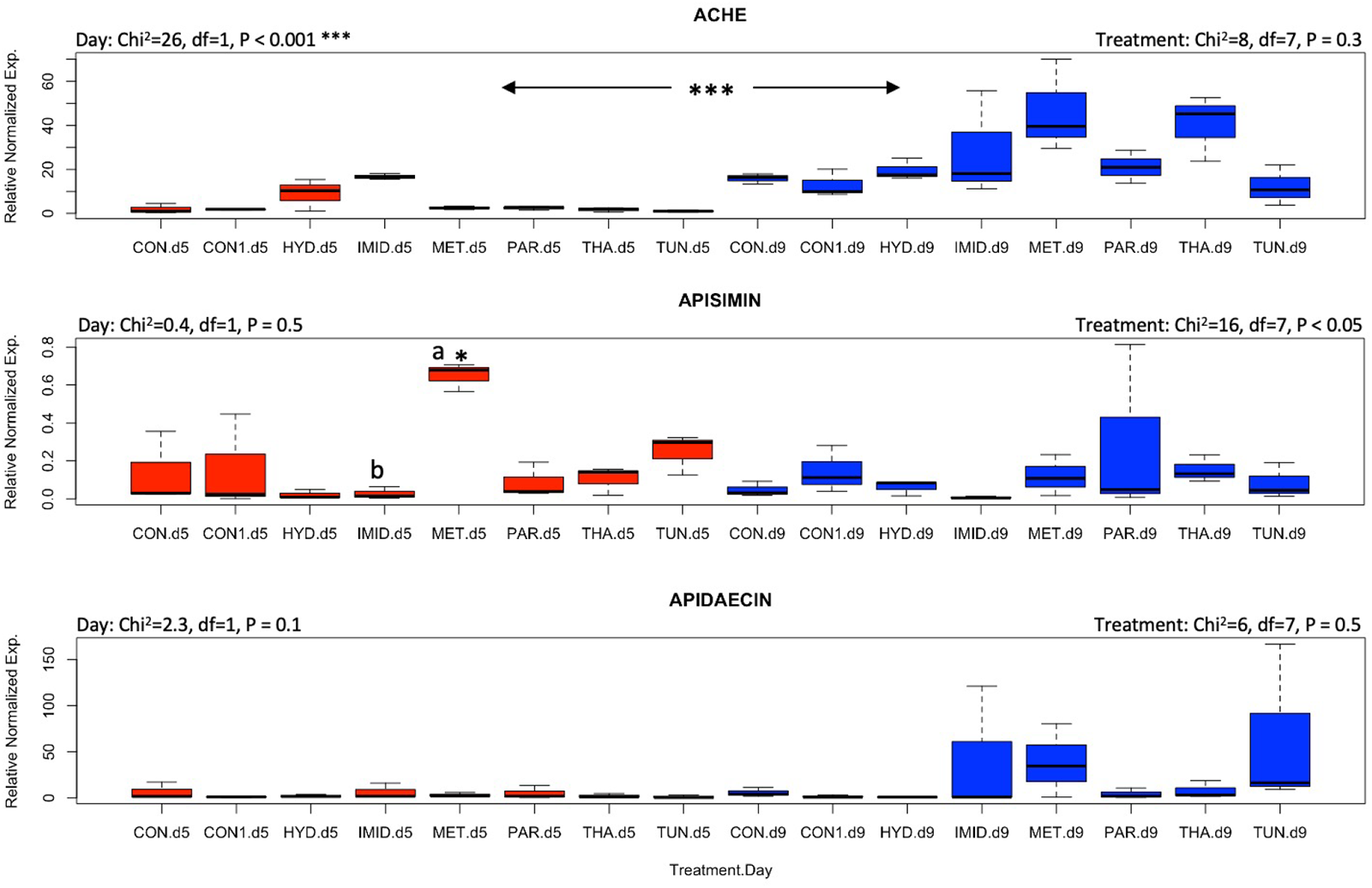
Overall gene expression of *AChE*, *apisimin* and *apidaecin* across eight treatments and days of exposure (Day 5: red boxplots and day 9: blue boxplots). Non-parametric Kruskal-Wallis test was conducted at a 95% confidential interval with three levels of significance (*p* < 0.05*, < 0.001**, < 0.001***) to determine statistical differences among treatments and dates. Boxplots with different alphabetical letters are statistically significant.

**Figure 4.**
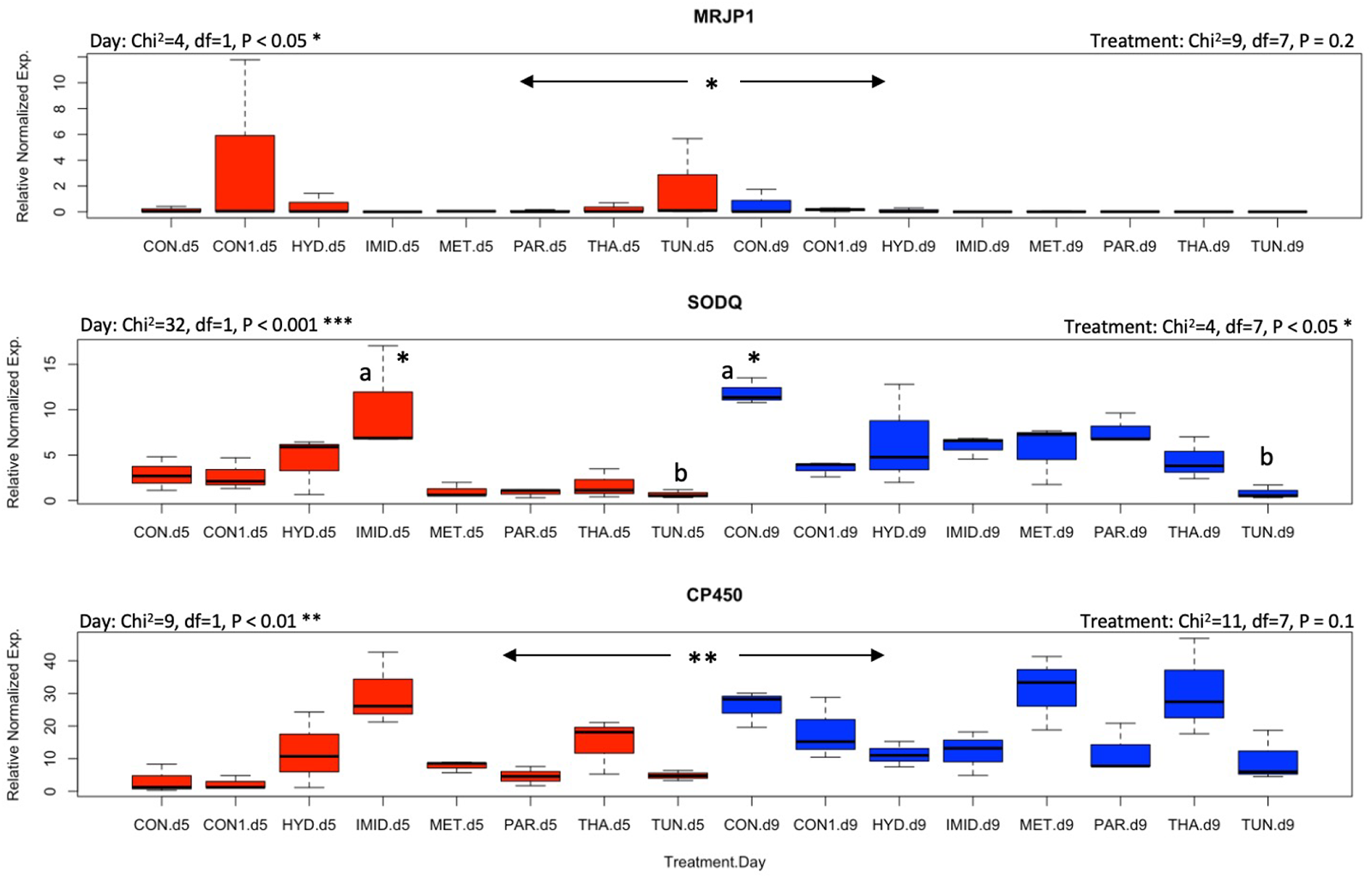
Overall gene expression of *mrjp1*, *Sodq* and *cp450* across eight treatments and studied dates (Day 5: red boxplots and day 9: blue boxplots). Non-parametric Kruskal-Wallis test was conducted at a 95% confidential interval with three levels of significance (*p* < 0.05*, < 0.001**, < 0.001***) to determine statistical differences among treatments and dates. Boxplots with different alphabetical letters are statistically significant.

**Figure 5.**
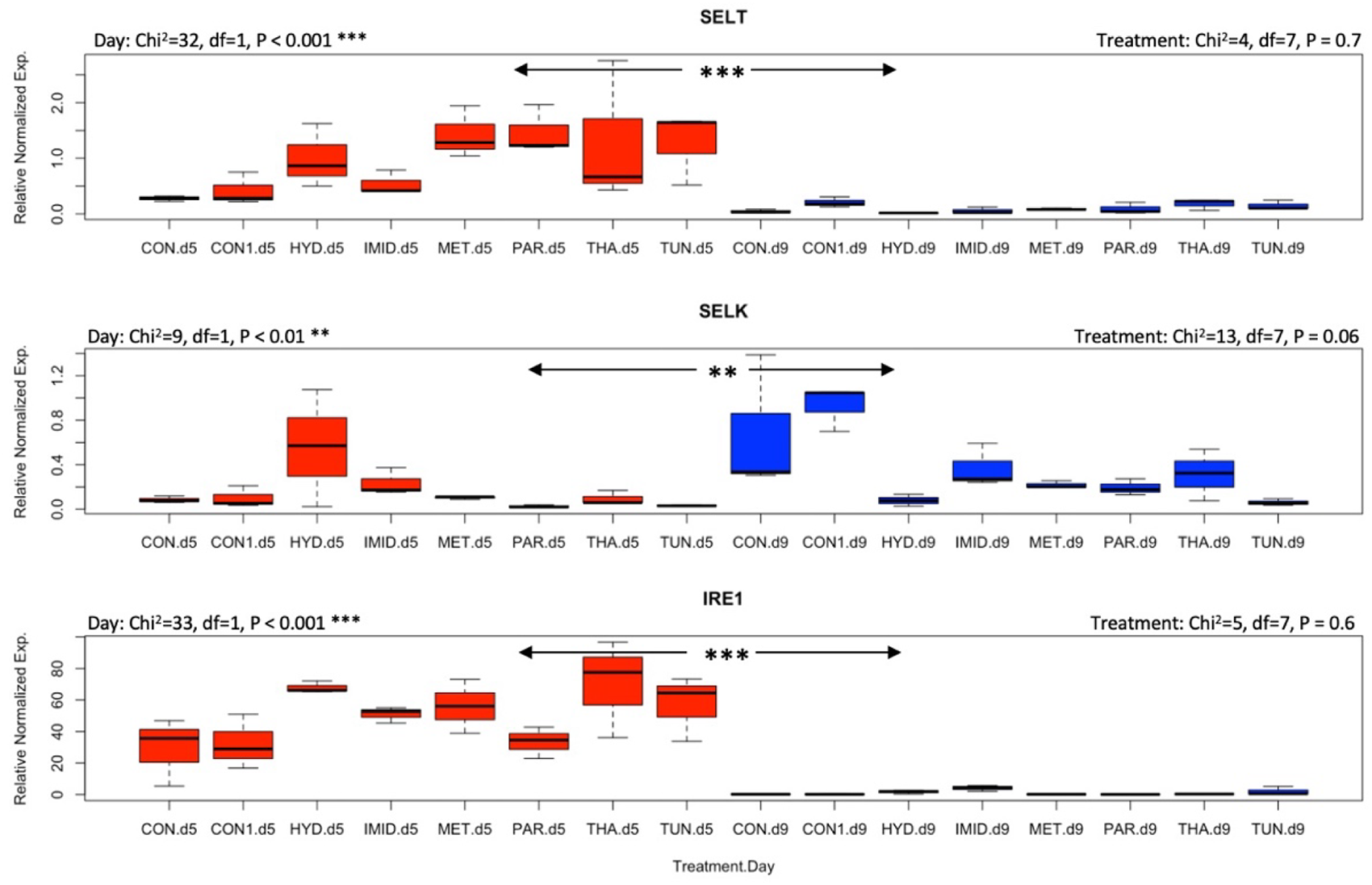
Overall gene expression of *SelT*, *SelK* and *Ire1* across eight treatments and studied dates (Day 5: red boxplots and day 9: blue boxplots). Non-parametric Kruskal-Wallis test was conducted at a 95% confidential interval with three levels of significance (*p* < 0.05*, < 0.001**, < 0.001***) to assess statistical differences among treatments and dates.

**Figure 6.**
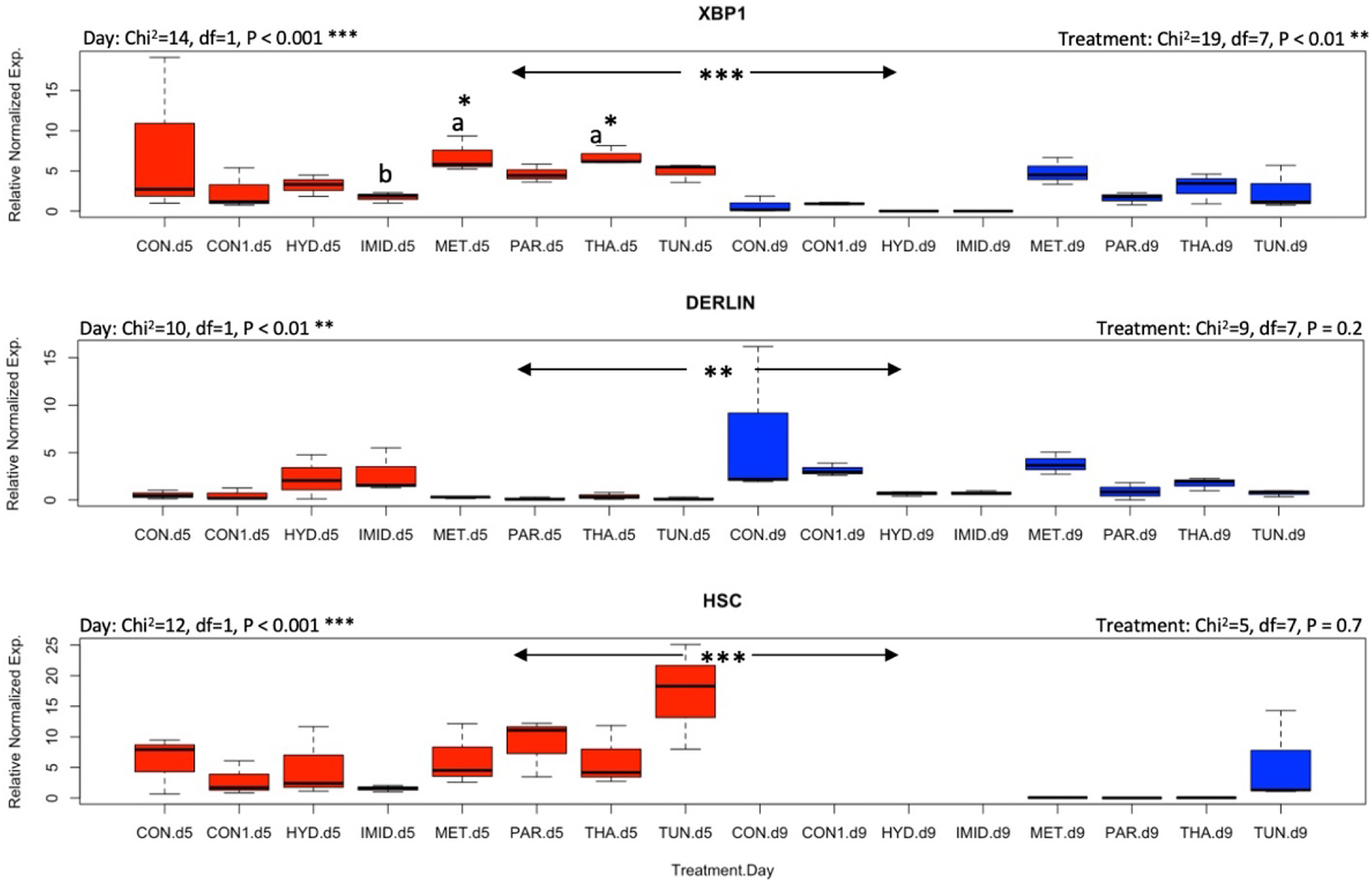
Overall gene expression of *Xbp1*, *Derl-1* and *Hsc70* across eight treatments and studied dates (Day 5: red boxplots and day 9: blue boxplots). Non-parametric Kruskal-Wallis test was conducted at a 95% confidential interval with three levels of significance (*p* < 0.05*, < 0.001**, < 0.001***) to assess statistical differences among treatments and dates. Boxplots with different alphabetical letters are statistically significant.

### 3. Gene Interaction and Correlation

The study of the overall gene regulation displayed as heatmaps revealed three different sets of genes: 1- Genes upregulated (*Ire1, AChE-2* and *Cp450*) and 2- Gene downregulated (*SelK, SelT, Apisimin, mrjp1*) and *3-* Genes with mix regulations (*Apideacin, Sodq, Hsc70, Xbp1, Derl-1*), Fig. 7a. The heatmap dendrogram distinguished four genes which exhibited the closest similarity in their overall regulations across treatments (CONT, PAR, MET, THA), Fig. 7a. However, this pattern of regulations was not constant and differed by dates. For instance, at day 5, the highest expressed gene among all treatments and genes was *Ire1,* while at day 9 upregulations were observed for *Apid*, *Sodq*, *AChE-2* and *Cp450* genes, Fig. 7b. The correlation analysis conducted on the overall and date-by-date gene regulations and treatments reveals significant (*p* < 0.01) positive correlations within each date only, Fig. 8. No significant negative correlations among genes were found at any time point.

**Figure 7.**
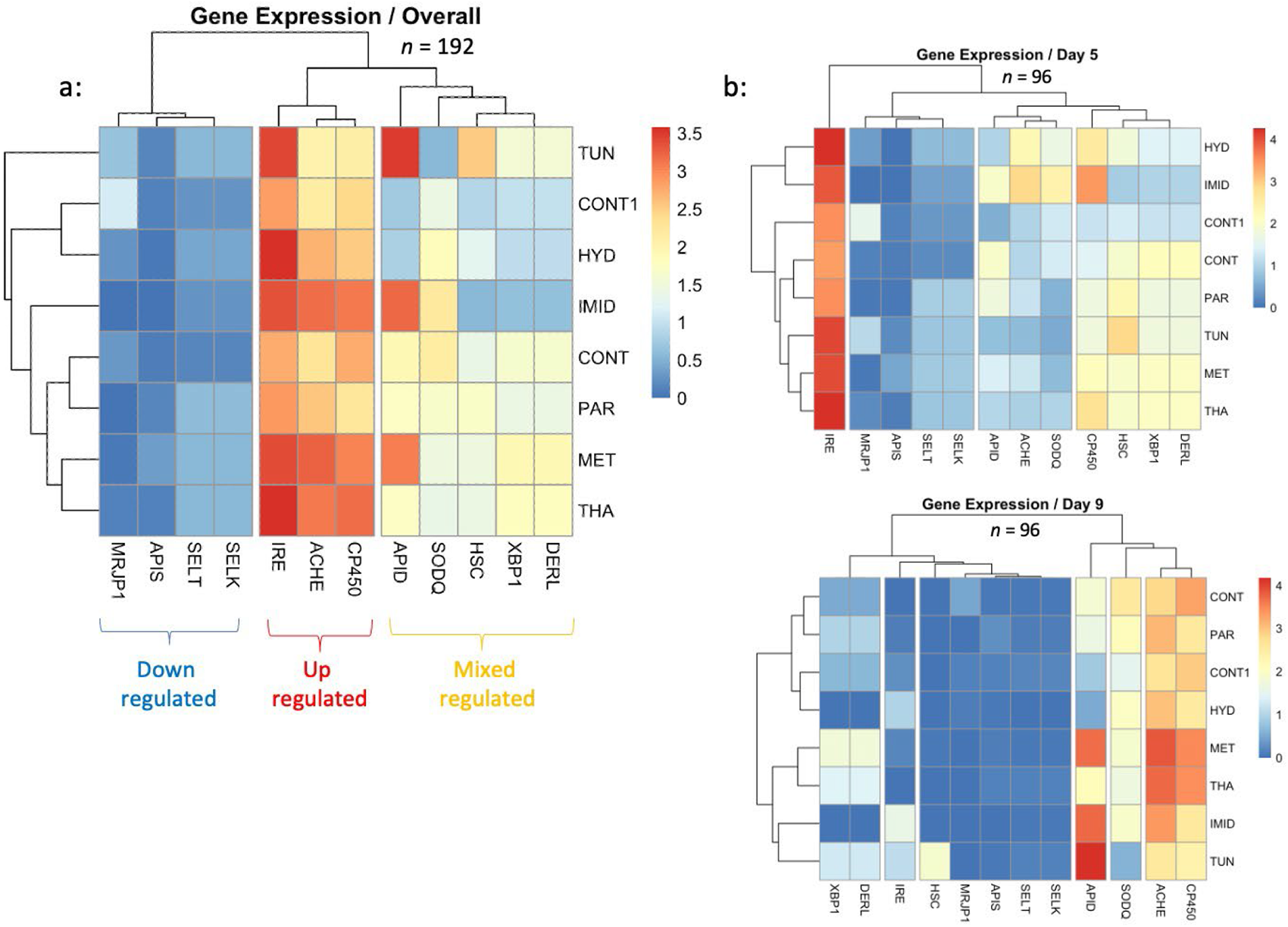
Heatmaps conducted on the overall regulation of the twelve studied genes in eight different treatments (a) as well as their regulations per date (b), (Day 5 and 9). Analysis of the overall gene regulation distinguished three major gene clusters showing upregulated (*Ire1*, *AChE- 2*, *Cp450*), downregulated (*SelK*, *SelT*, *Apis*, *mrjp1*), and mixed regulated genes (*Apis*, *Sodq*, *Hsc70*, *Xbp1, Derl-1*).

**Figure 8.**
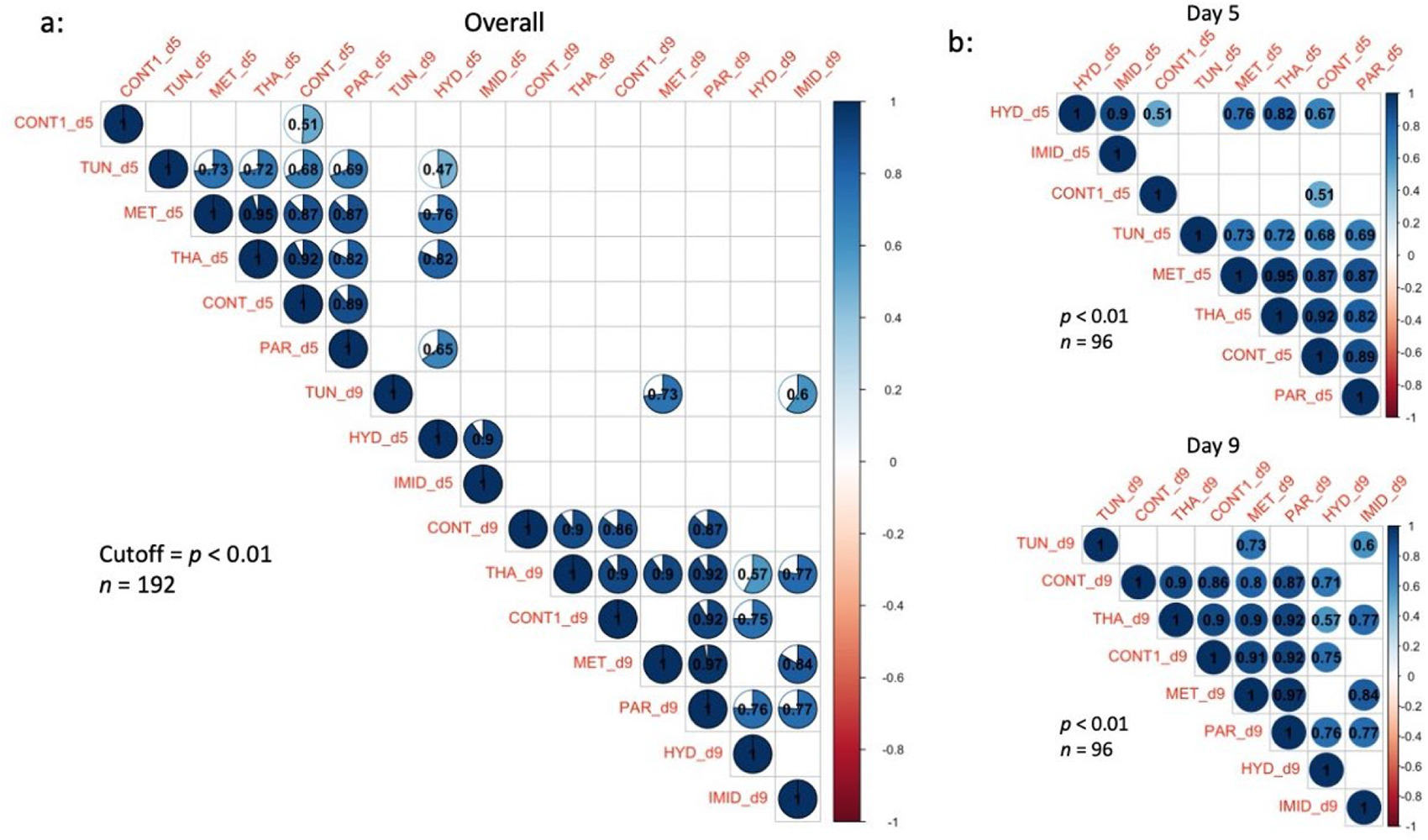
Correlation matrices of the gene regulation conducted on the eight studied treatments and displayed by overall (a) and date-by-date expressions (b). Correlation analysis was conducted at an intermediary level of significance (*p* < 0.01). Correlation R-values are given in each circle and blank squares represent non-significant correlations at the cutoff level of *p* < 0.01.

### 4. Principal Component Analysis

The PCA conducted on the regulation of all studied genes revealed that 40.7% of the variables were expressed on Dim1, 27.8% on Dim2 and 12.4% on Dim3, Fig. 9. According to the estimated number of clusters (K=3-4) and the visual distribution of the individual variables (Treatments), the PCA discriminates on Dim1 (40.7%) and Dim 2 (27.8%), four major groups which are: 1-(HYD and IMID), 2- (CONT and CONT1), 3- (MET and THA) and 4- (PAR and TUN), Fig. 9a. However, on Dim1 (40.7%) and Dim 3 (12.4%), treatments grouped into three clusters only: 1- (HYD, CONT1, IMID), 2- (CONT) and 3- (TUN, THA, PAR, MET), Fig. 9a.

**Figure 9.**
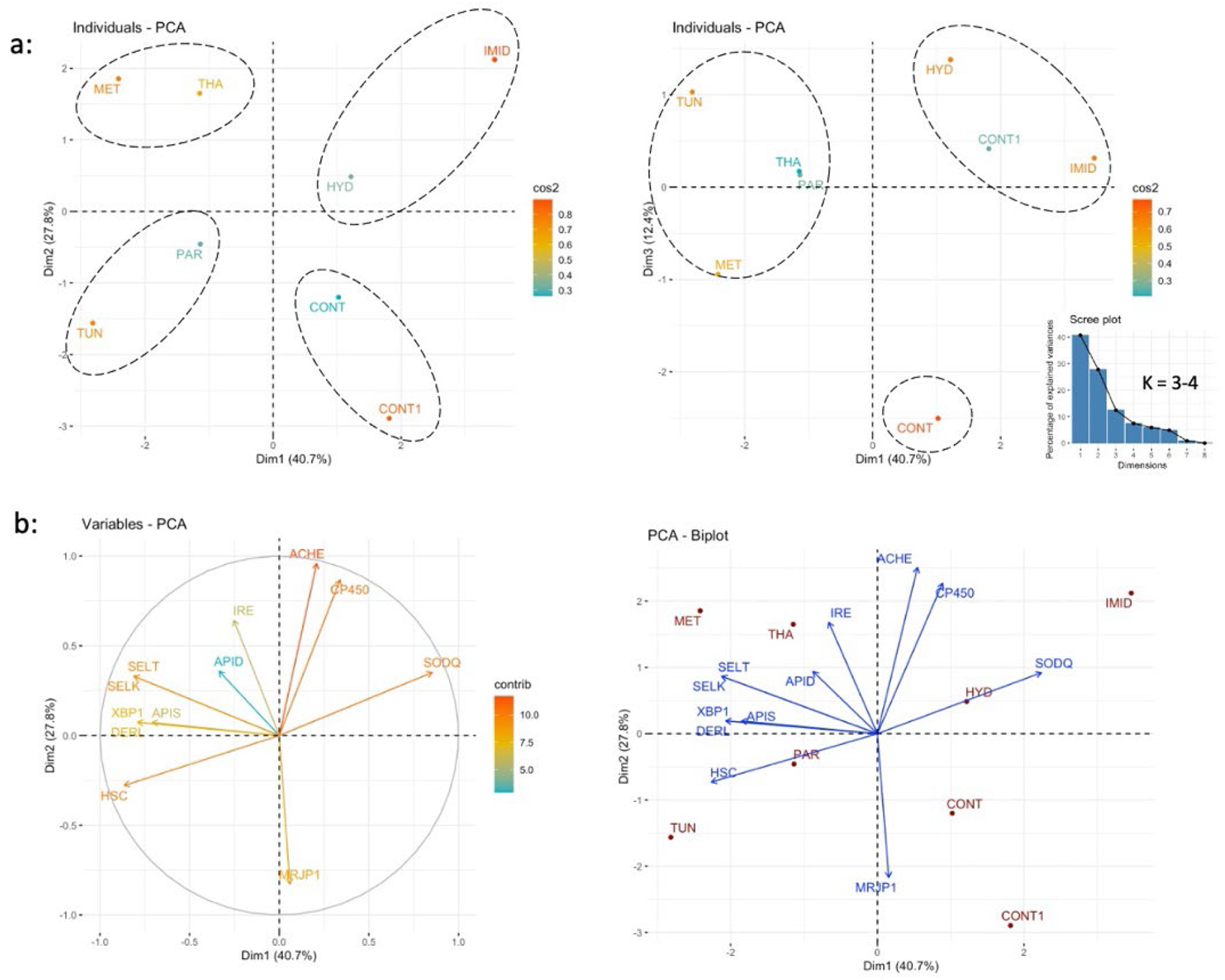
Principal Component Analysis (PCA) conducted on the overall regulation of the twelve studied genes in eight treatments. Percentages of individual variables expressed on components 1 and 2, 1 and 3 (a) graphically visualized in a 3-dimensional space. Expression and direction of each variable (genes) on Dim1 and 2 are given along with the PCA biplot of both expressed variables and treatments on Dim1 and 2 (b). The scree plot shows the mathematically calculated number of estimated groups (k).

Concerning the variable behavior (Regulation of the genes) regarding the treatments, the PCA displayed by “variables” showed two sets of genes with opposing function and regulation vis-à-vis the treatments: 1-(*cp450, Sodq)*, and 2- (*Ire1, Apis, SelT, SelK, Apid, Xbp1, Derl-1, Hsc70)*, while *mrjp1 and AChE-2* exhibited neutral regulation, Fig. 9b.

### 5. Oxidative Stress and Protein Damage

The protein carbonyl contents assay was conducted on caged bees sampled at day 9 of the treatment. The highest carbonyl contents were identified in bees fed paraquat and imidacloprid, which contained significantly (*p* < 0.001) higher protein damage than all other treatments. The control_H2O, metformin, thapsigargin, and tunicamycin did not statistically differ in their carbonyl contents, Fig. 10. The significantly lowest protein damage was identified in the control syrup containing PBS buffer (Control_PBS), Fig. 10.

**Figure 10.**
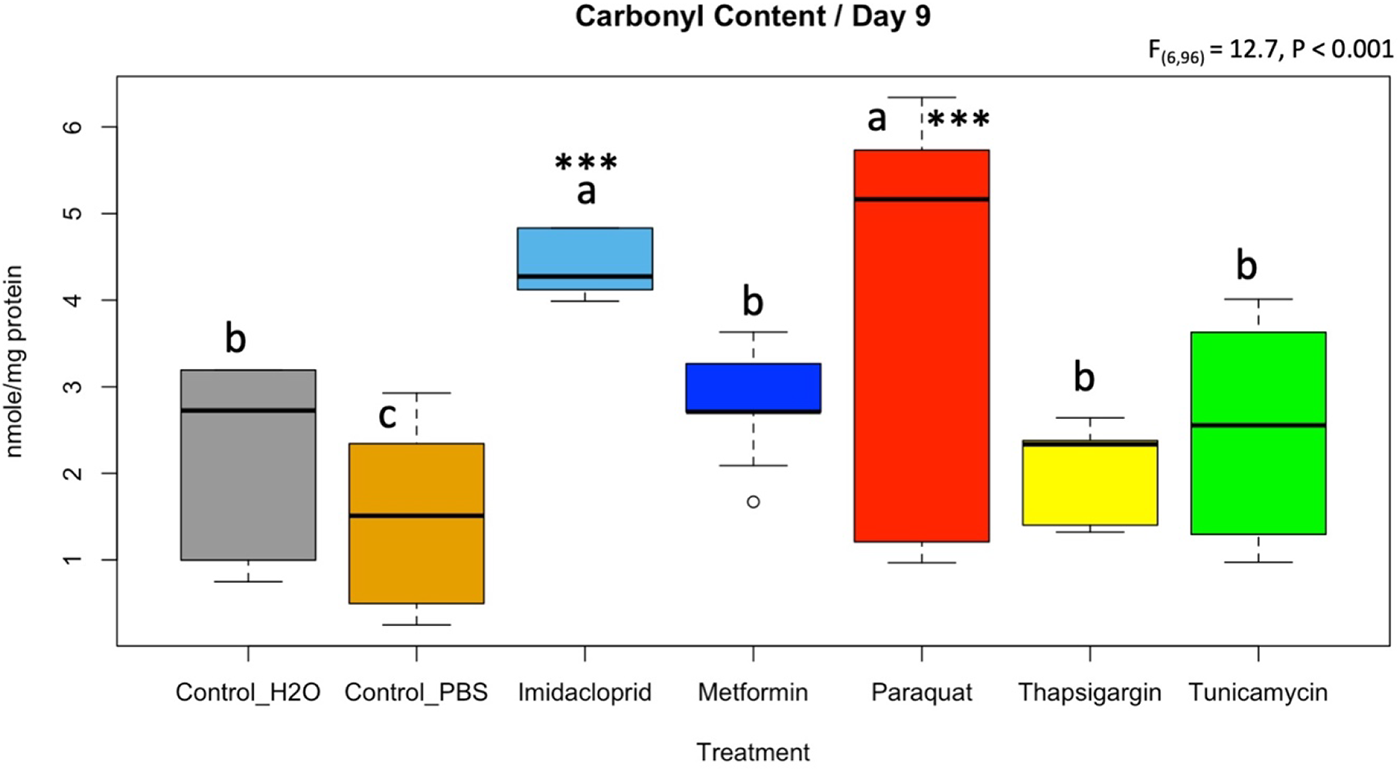
Level of protein damage quantified by Protein Carbonyl Content Assay on honey bee samples of Day 9 subjected to different treatments. ANOVA was conducted at a 95% confidential interval with three levels of significance (*p* < 0.05*, < 0.001**, < 0.001***) to assess statistical differences among treatments. Boxplots with different alphabetical letters are statistically significant.

## DISCUSSION

This study focused on the link between sublethal doses of pharmacological inducers and agricultural pesticides on honey bee gene regulation and oxidative stress. The eight treatments and controls used were a combination of known oxidative stress inducers in living organisms such as paraquat (herbicide), imidacloprid (neonicotinoid insecticide) and hydrogen peroxide (cellular byproduct of oxidative stress), as well as newly tested pharmacological compounds and antibiotics. Oxidative (or redox) stress reflects an imbalance between the systemic manifestation of reactive oxygen species (ROS) and a biological system’s ability to readily detoxify the reactive intermediates or repair the resulting damage (Pizzino *et al*. 2017). This imbalance can lead to ER stress, which occurs when proteins are not properly folded or conformed (Yamamoto AND ICHIKAWA 2019). For toxicity, our data showed that paraquat was the most chronically toxic molecule although it was administered at sublethal concentrations, Fig. 1. Paraquat catalyzes the formation of ROS through accepting electrons from photosystem I and transferring them to molecular oxygen (Kennedy *et al*. 2021). Our transcriptional results did not reveal specific gene response to alleviate the effect of paraquat on honey bees, however, the highest and most significant protein damage was recorded in honey bees when fed this treatment, Fig. 10. Moreover, the closest overall gene regulation similarity to paraquat was found in honey bees exposed to tunicamycin according to PCA analysis, Fig. 9a. Tunicamycin is an antibiotic which inhibits N-linked glycosylation inducing ER stress (Yamamoto AND ICHIKAWA 2019). Exposure to tunicamycin nonetheless produced only half (∼2.5 mole/mg) of the carbonyl content recorded for paraquat (∼5.1 mole/mg), Fig. 10. Despite clear refrain from consuming this antibiotic by honey bees compared to other molecules (*p* < 0.001, Fig. 2a), tunicamycin exhibited acute toxicity at day 8 (Fig. 1) with induced transcriptional response for *Hsc70* at day 9, Fig. 6. Heat shock 70-kDa protein plays important roles in normal cellular function and homeostasis. For instance, the Hsp70s function as molecular chaperones, assisting in protein synthesis, folding, assembly, trafficking between cellular compartments, and degradation) (Sarioglu-Bozkurt *et al*. 2022). It is conceivable that its higher regulation at day 5 alleviated the rate of protein damage identified at day 9 for tunicamycin which was one of the most toxic molecules tested, Fig. 1.

In honey bees, exposure to neonicotinoids has been well linked to elevated expressions of *AChE-2* (Badiou-Beneteau *et al*. 2012; Boily *et al*. 2013; Alburaki *et al*. 2015; Alburaki *et al*. 2023). Interestingly, our data confirmed previous findings related to exposure to imidacloprid in particular. Avoidance of sugar syrup tainted with imidacloprid (*p* < 0.001, Fig. 2), also recorded in this current study for tunicamycin and H_2_O_2_, was reported in previous investigations as well as significantly higher protein damage in caged bees exposed to this insecticide (Alburaki *et al*. 2019a; Alburaki *et al*. 2022). Expression of *AChE-2* was induced in bees exposed to imidacloprid at day 5 but remained statistically not significant compared to the control, Fig. 3. However, this gene significantly upregulated in all treatments at day 9 (Fig. 3), signaling potential occurrence of caging stress reported in a previous study (Alburaki *et al*. 2019a).

*Apisimin* was upregulated in a single occurrence in the case of bees exposed to metformin compared to honey bees exposed to imidacloprid, Fig. 3. Similar to *apidaecin*, both genes are described to play an important role in honey bee nutrition and are also antimicrobial peptides (AMPs) (Casteels *et al*. 1989; Srisuparbh *et al*. 2003; Shen *et al*. 2007). *Apidaecin* showed no regulation of any kind in our study, Fig. 3. Hydrogen peroxide which works by producing destructive hydroxyl free radicals that can attack membrane lipids, DNA, and other essential cell components (Brudzynski 2020), was avoided by bees (Fig. 2a) and led to significant early toxicity when administered at 4,000 ppb, Fig. 1. The closest similarity of the overall gene response to H_2_O_2_ according to the PCA is that of imidacloprid, Fig. 9a. Furthermore, our transcriptional data revealed crucial data related to genes potentially linked to the behavioral caste development in honey bees, Figs 4,5,6. A set of genes was significantly regulated based on the factor “Time” or “Age of the honey bees” irrespective of the treatments administered. This set of genes includes *mrjp1*, *Cp450*, *SelT*, *SelK*, *Derl-1*, *Ire1* and *Hsc70*. The first gene (*mrjp1*), which is well known to downregulate in foragers compared to nurse bees, has already been described as a physiological marker for behavioral development (Corona *et al*. 2023). The other genes require further investigations pertaining to their age-related regulations in honey bees.

In conclusion, the treatments of tunicamycin, hydrogen peroxide, and imidacloprid showed signs of disinclination in post-ingestion as well as a pharmacological enhancement in bee survivorship compared to the honey bees fed sugar syrup control. While a few antioxidant genes were significantly regulated in regard to the different treatments, our results reveal age-related regulation of other major genes with significant inter-gene positive correlations. Lastly, significant protein damage in the honey bee was observed in the treatments of paraquat and imidacloprid when administered *ad libitum* for 11 days.

## Author contributions

Study conception and design: SK, FT, JA, MA; Fund acquisition: SK, JA, MA, Data collection: FT, MA; Data analysis: MA, MG, SK, wrote the manuscript: FT, MA, revised it: SK, JA. All authors revised and approved the final version of this manuscript.

## Data Availability

All data related to this study are included in this manuscript and its supplementary materials.

## Additional Information

The authors declare no conflict of interest to report. This research was funded by a USDA ARS cooperative agreement # 58-6066-9-041 (Southern Horticultural Lab, Poplarville, MS); the USDA-ARS Bee Research Laboratory in Beltsville, MD, USA; USDA NIFA award #2023-67014- 39916. We thank Mississippi INBRE, supported by the NIH NIGMS (P20GM103476) for using the instrumentation core. The funders played no role in the study design, data collection, analysis, publication, decision or manuscript preparation.

